# Natural *SEL1L* variants modify ERAD, proteasome function, and survival in a *Drosophila* model of NGLY1 deficiency

**DOI:** 10.1101/2025.03.06.641902

**Authors:** Travis K. Tu’ifua, Clement Y. Chow

**Affiliations:** Department of Human Genetics, University of Utah School of Medicine, Salt Lake City, Utah, United States of America

## Abstract

N-glycanase 1 (NGLY1) deficiency is an ultra-rare disease caused by autosomal recessive loss-of-function mutations in the *NGLY1* gene. NGLY1 removes N-linked glycans from glycoproteins in the cytoplasm and is thought to help clear misfolded proteins from the endoplasmic reticulum (ER) through the ER associated degradation (ERAD) pathway. Despite this, the physiological significance of NGLY1 in ERAD is not understood. The best characterized substrate of NGLY1 is NRF1, a transcription factor that upregulates proteasome expression and the proteasome bounce-back response. We previously performed a genetic modifier screen using a *Drosophila* model of NGLY1 deficiency and identified potential modifiers that alter the lethality of the model. We identified two protein-coding variants in *Hrd3*/*SEL1L*: *S780P* and *Δ806-809*. Both variants are localized to the SEL1L cytoplasmic tail, an uncharacterized domain. SEL1L is a component of the ERAD complex that retrotranslocates misfolded proteins from the ER to the cytoplasm for degradation. We used CRISPR to generate fly lines carrying these *SEL1L* variants in a common genetic background and tested them with our model of NGLY1 deficiency. Validating our previous screen, the *SEL1L^P780^* and *SEL1L^Δ806-809^* variants increase the survival of the NGLY1 deficiency model, compared to the *SEL1L^S780^* variant. To determine how these *SEL1L* variants were modifying lethality in NGLY1 deficiency, we interrogated the ERAD and NRF1 signaling pathways. We found that the *SEL1L^P780^* and *SEL1L^Δ806-809^* variants improve ERAD function in an NGLY1- dependent manner, further implicating NGLY1 in general ERAD function. We also found that these variants protect against changes in larval size and survival caused by proteasome inhibition in heterozygous *NGLY1* null flies. These results provide new insights into the role of SEL1L in the disease pathogenesis of NGLY1 deficiency. *SEL1L* is a strong candidate modifier gene in patients, where variability in presentation is common.

**AUTHOR’S SUMMARY:** NGLY1 deficiency is a debilitating rare genetic disorder. There are currently no treatment options for NGLY1 deficiency and NGLY1 biology remains poorly understood. We previously performed a genetic modifier screen in a *Drosophila* model of NGLY1 deficiency and identified a number of candidate modifier genes that impacted the survival of our model. Modifier genes can help reveal NGLY1 biology and NGLY1 deficiency disease pathogenesis. In this study, we follow-up on two natural protein-coding variants of *Hrd3* (the fly version of the human gene, *SEL1L*) that increased the survival of our NGLY1 deficiency model. SEL1L is a critical component of an important quality control pathway called the endoplasmic reticulum associated degradation (ERAD) pathway. We discovered that these *SEL1L* variants enhance ERAD and modify NGLY1 deficiency sensitivity to proteasome inhibition. This study confirms *SEL1L* as an important modifier gene of NGLY1 deficiency. Further study of the ERAD and proteasome degradation pathways may reveal additional candidate modifier genes of NGLY1 deficiency and potential targets for therapeutic development.

## INTRODUCTION

N-glycanase 1 (NGLY1) deficiency is an ultra-rare disease and the first identified congenital disorder of deglycosylation (CDDG). The disease is caused by autosomal recessive loss-of-function mutations in the *NGLY1* gene (1,2). NGLY1 is a cytosolic deglycosylating enzyme that removes N-linked glycans from proteins. NGLY1 is thought to be a component of the endoplasmic reticulum (ER) associated degradation (ERAD) pathway, an important cellular quality control mechanism which removes misfolded proteins from the ER to the cytosol for degradation (3,4). However, the loss of NGLY1 shows little effect on ERAD and does not prevent the degradation of misfolded proteins (5–7). Therefore, despite its known function as a deglycosylating enzyme, the physiological significance of NGLY1 and disease pathogenesis remains poorly understood.

NGLY1 deficiency is marked by extensive phenotypic heterogeneity, even among patients with identical *NGLY1* mutations (2,8), suggesting the presence of genetic modifiers. In a previous genetic screen, we identified 61 potential modifier genes that were associated with changes in survival in our *Drosophila* model of NGLY1 deficiency (9). From this screen, our top hit was *Ncc69* (*Drosophila* ortholog for human *NKCC1/2*), which encodes for a conserved ion transporter and we showed that it is both a substrate of NGLY1 and a modifier of NGLY1 deficiency (9). Another interesting candidate modifier gene we identified in the screen was *Hrd3* (hereon referred to by the human ortholog *SEL1L*). Through a genome-wide association study (GWAS), we identified a natural missense variant in *SEL1L* that was associated with increased survival in the *Drosophila* NGLY1 deficiency model (9). This *SEL1L* variant is a substitution of serine 780 (*SEL1L^S780^*) for a proline (*SEL1L^P780^*). Additionally, we identified a private protein-coding deletion in the strain showing near complete rescue of NGLY1 deficiency lethality. This second variant is a deletion of amino acids 806 to 809 (*SEL1L^Δ806-809^*). Both variants are 26 amino acids apart and are located in the cytoplasmic tail of SEL1L.

SEL1L is a single-pass ER membrane protein and a critical, well-established component of ERAD. ERAD functions alongside other quality control mechanisms such as the unfolded protein response (UPR) and autophagy to maintain ER homeostasis and prevent ER stress (10–12). The SEL1L-Hrd1 ERAD complex is the most conserved branch of ERAD from yeast to humans and translocates misfolded proteins from the ER to the cytosol for proteasomal degradation (13–15). The luminal domain of SEL1L assists in the recognition of ERAD substrates in the ER lumen and the transmembrane domain helps move proteins through the ER membrane to the cytosol (16). The cytoplasmic tail of SEL1L is a highly disordered region across species and its function is unknown.

In previous studies, both *SEL1L* and *NGLY1* were identified as genetic modifiers of NRF1, a transcription factor responsible for the proteasome bounce-back response (17,18). NRF1 is co-translated and glycosylated in the ER before ERAD machinery translocates NRF1 to the cytosol (19). NGLY1 deglycosylation and DDI1/2 protease cleavage activate NRF1. Deglycosylation of N-linked glycans by NGLY1 results in the deamidation of asparagine to aspartate residues. This amino acid editing is necessary and sufficient for NRF1 activation, localization, and function (17,18). The protease DDI1/2 cleaves NRF1 to release the activated protein from its ER tether (17,18,20). Under healthy, homeostatic conditions, activated NRF1 is constitutively degraded by the proteasome. Under conditions of proteasomal stress, however, NRF1 is not degraded, accumulates in the cytosol, and is transported to the nucleus where it acts as a transcription factor and upregulates genes that increase proteasome function, including proteasome subunit genes (17,18,21). This activation of NRF1 is known as the proteasome bounce-back response. NGLY1 and ERAD machinery are necessary for NRF1 activation and the loss of either prevents the proteasome bounce-back response (17,18).

In this study, we characterized the functional consequences of the two new *SEL1L* variants identified in our NGLY1 deficiency genetic screen. We placed the *SEL1L* variants on an isogenic background to test the effects each variant has on both SEL1L and NGLY1. The *SEL1L* variants increase survival in our NGLY1 deficiency model, validating the observations from the modifier screen. The *SEL1L* variants also enhance ERAD in an NGLY1-dependent manner and provide a protective fitness advantage during proteasome inhibition. Our results suggest that *SEL1L* is a modifier of *NGLY1* and that interactions between *SEL1L*, *NGLY1*, and *NRF1* underly these observed changes in fitness. These genetic interactions are potential targets for NGLY1 deficiency treatment.

## RESULTS

### *SEL1L^P780^* and *SEL1L^Δ806-809^* variants increase survival of NGLY1 deficiency model

In a previous study, we crossed our NGLY1 deficiency *Drosophila* model, which uses the *GAL4/UAS* system to ubiquitously express RNAi against *NGLY1*, with nearly 200 strains of the *Drosophila* Genetic Reference Panel (DGRP) (9,22). On a standard laboratory background, the NGLY1 deficiency model has ∼30% survival to adulthood (7,9). In the DGRP strains, the NGLY1 deficiency model showed survival ranging from 0 to 100%, indicating that lethality is highly modifiable by genetic background. We performed a GWAS to identify candidate modifier genes associated with increased survival. One of the top associated variants was the S780P missense variant in *SEL1L*. The *SEL1L^P780^* minor allele was associated with increased survival of the NGLY1 deficiency model, compared to the common *SEL1L^S780^* allele. We also discovered that the DGRP strain with nearly 100% survival in the screen (DGRP strain 379) harbored a private *SEL1L* variant, resulting in the deletion of amino acids 806-809 (*SEL1L^Δ806-809^*). This strain also carries the more common *S780* allele. *SEL1L^Δ806-809^* was not formally identified through the GWAS because it is a private variant in a single DGRP strain. Because these two variants were both in the cytoplasmic tail of SEL1L, a functionally uncharacterized region of the protein, we sought to understand how these variants were affecting survival of the NGLY1 deficiency model. We used CRISPR to place each of the *SEL1L* variants (Fig 1A) onto the same isogenetic background, creating three strains that are homozygous for *SEL1L^S780^*, *SEL1L^P780^,* or *SEL1L^Δ806-809^*. This allowed us to test for phenotypic differences specific to each *SEL1L* variant.

**Fig 1.**
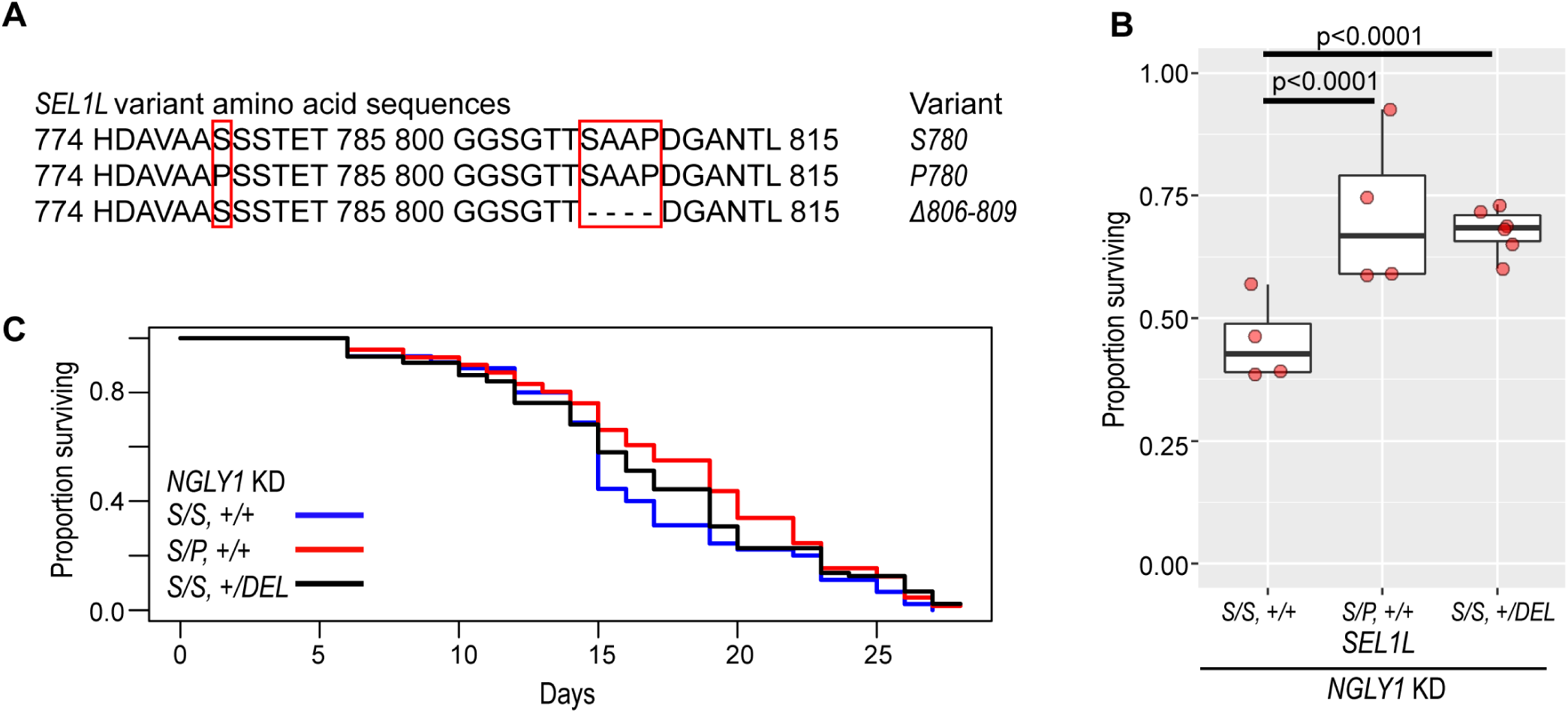
*SEL1L* variants increase NGLY1 deficiency survival. (A) Amino acid sequences for the three *SEL1L* variant alleles. Red boxes highlight differences between alleles. (B) *SEL1L^(S/P, +/+)^* and *SEL1L^(S/S, +/DEL)^* genotypes significantly increase the proportion surviving (∼70%) with *NGLY1* knockdown when compared to *SEL1L^(S/S, +/+)^* (45%, p<0.0001). Chi-squared test. (C) Lifespan of *NGLY1* KD flies shows no significant difference in survival between *SEL1L* genotypes. Cox proportional hazard regression analysis

In our original screen, we crossed a strain carrying both a GAL4 and *NGLY1 RNAi* transgene with strains of the DGRP. Lethality was scored based on survival of the F1 flies, which had half of their genomes coming from the *NGLY1* RNAi strain and half from the different DGRP strains. Because the *NGLY1* RNAi strain is homozygous for the common *SEL1L^S780^* allele, any new *SEL1L* variant introduced by the DGRP strain in the F1 generation was heterozygous with the *SEL1L^S780^* allele. Therefore, the relevant *SEL1L* genotypes from our screen are: *SEL1L^(S/S, +/+)^*, *SEL1L^(S/P, +/+)^*, or *SEL1L^(S/S, +/DEL)^*. We focus our analyses on these three *SEL1L* genotypes throughout this study.

Based on the original DGRP screen, we expected that the *SEL1L^(S/P, +/+)^* and *SEL1L^(S/S, +/DEL)^* genotypes would increase survival of the NGLY1 model, compared to the *SEL1L^(S/S, +/+)^* genotype. To validate the results of the screen, we crossed the same *NGLY1* RNAi strain used in the modifier screen with each of our new *SEL1L* variant CRISPR strains to generate flies that have the exact *SEL1L* genotypes from the screen. The proportion surviving was determined in the same manner as the original screen, by dividing the number of *NGLY1* knockdown flies by the largest balancer class in its cross (9). There was significantly increased survival of NGLY1 knockdown flies with the *SEL1L^(S/P, +/+)^* and *SEL1L^(S/S, +/DEL)^* genotypes compared to the *SEL1L^(S/S, +/+)^* genotype (Fig 1B and S1 Data). The *SEL1L^(S/S, +/+)^* genotype had a 45% proportion surviving compared to ∼70% in the *SEL1L^(S/P, +/+)^* (p<0.001) and *SEL1L^(S/S, +/DEL)^* (p<0.001) genotypes. This result nicely replicates our previous genetic screen that showed increased survival in the DGRP lines with these particular *SEL1L* variants and suggests that the *SEL1L^P780^* and *SEL1L^Δ806-809^* alleles are protective against NGLY1 deficiency. We also evaluated the lifespan of these surviving NGLY1 knockdown flies, but found no significant differences in survival between the *SEL1L^(S/P, +/+)^* and *SEL1L^(S/S, +/DEL)^* and the *SEL1L^(S/S, +/+)^* genotype (Fig 1C and S1 Data), suggesting that the interaction occurs during development.

### *SEL1L* variants impact sensitivity to proteasome inhibition in *NGLY1^+/-^* flies

We next interrogated pathways that involve both SEL1L and NGLY1 to understand how these *SEL1L* variants are protecting against NGLY1 deficiency. Both *NGLY1* and *SEL1L* were previously identified as modifier genes of *NRF1*, which encodes for an important transcription factor that upregulates proteasome genes in response to proteasomal stress (17,18). The importance of NGLY1 in NRF1 function is well established. NGLY1 mutants have reduced proteasome function and are exquisitely sensitive to proteasome stress because NRF1 is not processed (17,18,23,24). Previous studies have shown that heterozygous *NGLY1* null larvae, which are otherwise normal, are sensitive to proteasome inhibition, leading to larval size defects (23,24). Although *SEL1L* was identified as a genetic modifier of NRF1, its role in NRF1 signaling has not been determined. We hypothesized that these *SEL1L* variants would affect NRF1 signaling and modify phenotypes in an NGLY1 deficiency model.

We tested whether the *SEL1L* variants affect proteasome sensitivity in NGLY deficient *Drosophila* using the proteasome inhibitor bortezomib (BTZ). Homozygous *NGLY1* null *Drosophila* are embryonic lethal, but heterozygous *NGLY1* null flies are phenotypically normal when unchallenged. We used heterozygous *NGLY1* null flies as a model of NGLY1 deficiency because of their known increased sensitivity to proteasome inhibition (23,24). Because the *NGLY1* null strain also carries the common *SEL1L^S780^* allele, when we cross this strain with our CRISPR generated *SEL1L* variant strains, we generate heterozygous *NGLY1* null flies with the same *SEL1L* genotypes to what we tested in the *NGLY1* knockdown model: *SEL1L^(S/S, +/+)^*, *SEL1L*^(*S/P*,+/+)^, or SEL1L*^(S/S, +/DEL)^*.

Heterozygous *NGLY1* null *Drosophila* larvae develop smaller when exposed to proteasome inhibition, compared to *NGLY1* wildtype and DMSO-treated heterozygous *NGLY1* null controls (23,24). In previous studies, heterozygous *NGLY1* null larvae, when exposed to 5μM BTZ, are significantly smaller than DMSO-treated heterozygous *NGLY1* null larvae (24). We observed an equally strong decrease in larval size with the treatment of 5μM bortezomib in all heterozygous *NGLY1* null larvae compared to DMSO controls, regardless of *SEL1L* genotype (Fig 2A and S2 Data). *SEL1L* genotype does not impact the size defects induced by 5μM BTZ in *NGLY1* heterozygous null larvae.

**Fig 2.**
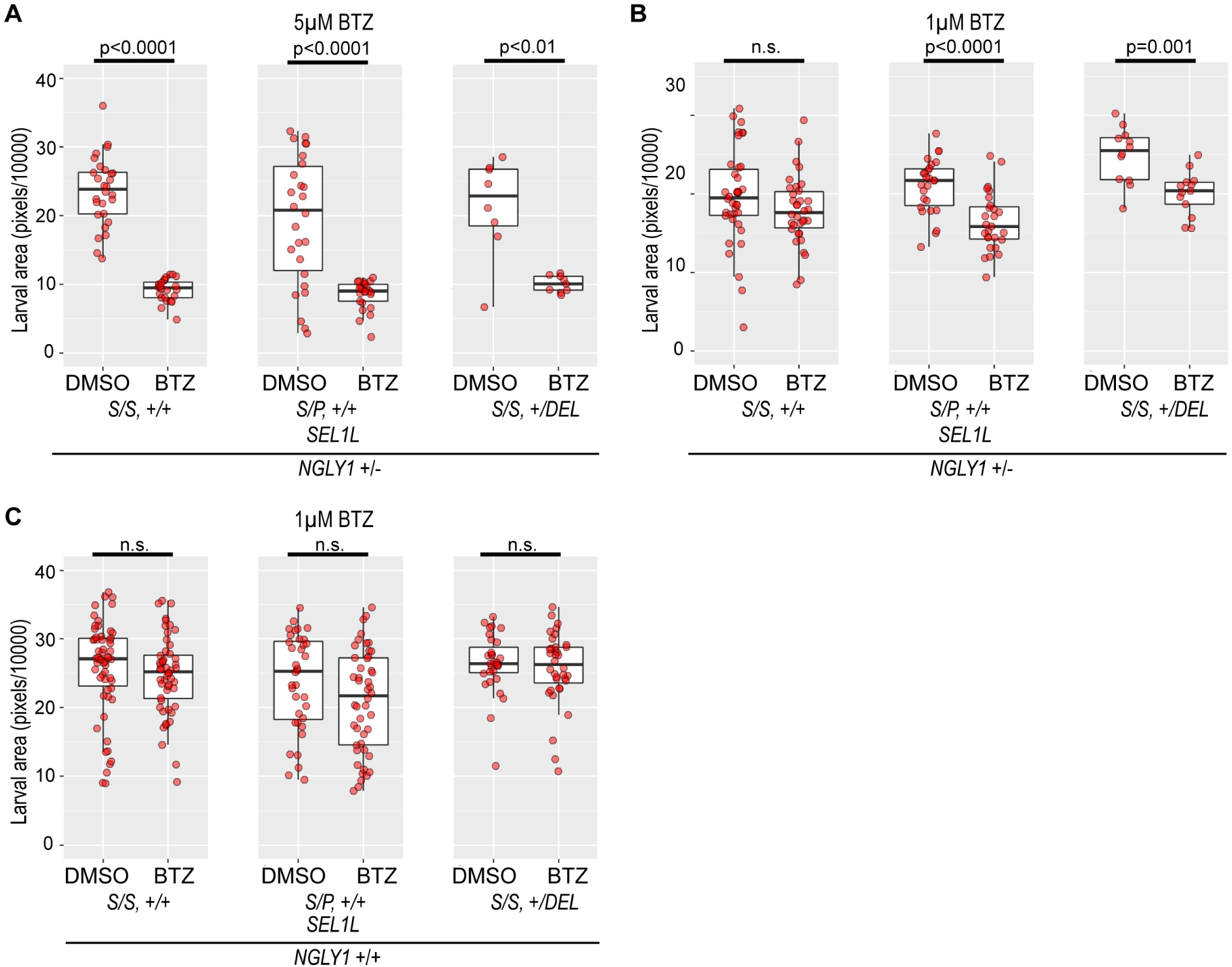
*SEL1L* variants affect proteasome inhibition sensitivity. (A) *NGLY1 +/-* larvae are smaller when treated with 5μM BTZ, but there is no effect of *SEL1L* genotype (Larval size on DMSO: *SEL1L^(S/S, +/+)^* 23.59 ± 4.97, *SEL1L^(S/P, +/+)^* 19.37 ± 9.16, and *SEL1L^(S/S, +/DEL)^* 21.32 ± 6.73. Decreased larval size on BTZ: *SEL1L^(S/S, +/+)^* 9.19 ± 1.61, p<0.0001; *SEL1L^(S/P, +/+)^* 8.55 ± 2.04, p<0.0001; and *SEL1L^(S/S, +/DEL)^* 10.07 ± 1.14, p<0.01). (B) When treated with 1μM BTZ, the *NGLY1 +/-* larvae show a *SEL1L* genotype-dependent decrease in size. While *SEL1L^(S/S, +/+)^* showed no change between DMSO (19.65 ± 5.98) and BTZ (17.88 ± 4.32), *SEL1L^(S/P, +/+)^* (DMSO 21.03 ± 3.42, BTZ 16.45 ± 3.65, p<0.0001) and *SEL1L^(S/S, +/DEL)^* (DMSO 24.92 ± 3.39, BTZ 20.08 ± 2.69, p=0.001) genotypes were smaller when treated with BTZ compared to DMSO treated larvae. (C) *NGLY1 WT* larvae show no significant decrease in size with 1μM BTZ treatment, regardless of *SEL1L* genotype (Larval size on DMSO: *SEL1L^(S/S, +/+)^* 25.75 ± 6.91, *SEL1L^(S/P, +/+)^* 23.74 ± 6.94, and *SEL1L^(S/S, +/DEL)^* 26.42 ± 4.29; larval size on BTZ: *SEL1L^(S/S, +/+)^* 24.75 ± 5.63, *SEL1L^(S/P, +/+)^* 20.96 ± 7.38, and *SEL1L^(S/S, +/DEL)^* 25.70 ± 5.23).

To determine whether there might be more subtle effects, we treated the heterozygous *NGLY1* null larvae with a lower concentration of 1μM BTZ. The *SEL1L^(S/S, +/+)^* larvae showed no significant size differences between BTZ and DMSO treatments. However, *SEL1L^(S/P, +/+)^* (p<0.0001) and *SEL1L^(S/S, +/DEL)^* (p= 0.001) larvae had significant decreases in larval size with 1μM BTZ treatment compared to DMSO (Fig 2B and S2 Data). This indicates that larval size in heterozygous *NGLY1* null larvae carrying the *SEL1L^(S/P, +/+)^* and *SEL1L^(S/S, +/DEL)^* genotypes are particularly sensitive to proteasome inhibition compared to *SEL1L^(S/S, +/+)^*. On an NGLY1 wildtype background, the different *SEL1L* genotypes showed no larval size changes with the 1μM BTZ treatment (Fig 2C and S2 Data).

### *SEL1L* variants increase survival of *NGLY1^+/-^* flies in response to proteasome inhibition

To further examine the impact of the *SEL1L* variants on NRF1 signaling, we tested other phenotypes affected by NGLY1 deficiency and proteasome inhibition. When heterozygous *NGLY1* null larvae were treated with 1μM BTZ, we observed a *SEL1L* variant specific effect on survival through eclosion to adulthood. The *SEL1L^(S/P, +/+)^* and the *SEL1L^(S/S, +/DEL)^* larvae showed higher survival at 83% (p<0.0001) and 96% (p<0.0001), respectively, compared to the *SEL1L^(S/S, +/+)^* flies at 59% survival (Fig 3A and S3 Data). *SEL1L^(S/S, +/+)^* larvae treated with bortezomib showed high rates of non-eclosed and partially eclosed flies. The improved eclosion and survival rates of the *SEL1L^(S/P, +/+)^* and the *SEL1L^(S/S, +/DEL)^* genotypes suggests a protective effect of the *SEL1L^P780^* and *SEL1L^Δ806-809^* variants against NGLY1 deficiency and proteasome inhibition during larval development. Importantly, when *NGLY1* wildtype larvae were raised on 1μM BTZ, we observed no lethality and nearly 100% eclosion of flies, regardless of *SEL1L* genotype, demonstrating that lethality to BTZ is NGLY1-dependent (S1 Fig and S3 Data). *SEL1L^(S/P, +/+)^* and *SEL1L^(S/S, +/DEL)^* genotypes provide near complete rescue of the heterozygous *NGLY1* null larvae lethality induced by 1μM BTZ.

**Fig 3.**
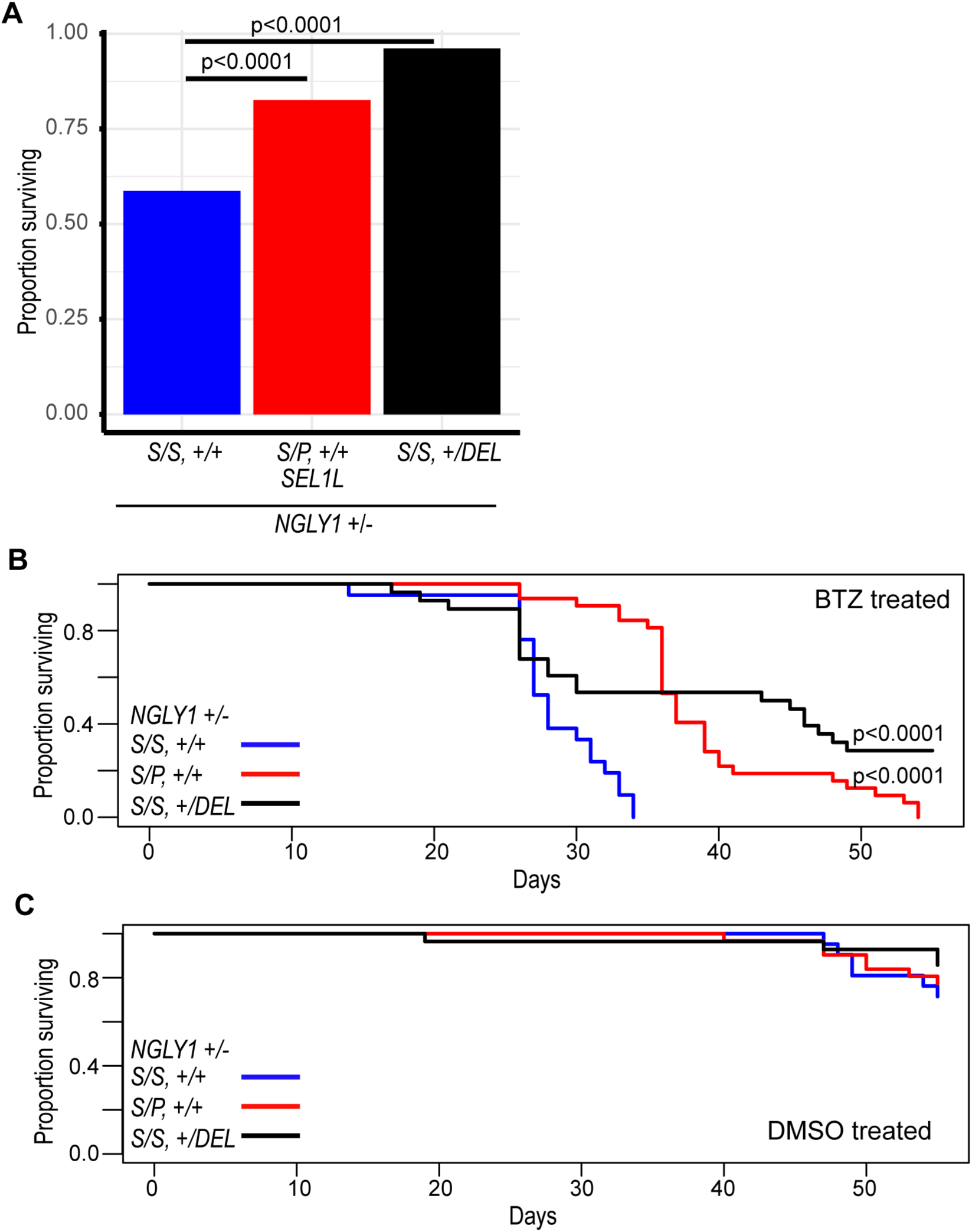
*SEL1L* variants increase survival under proteasome inhibition. (A) When treated with 1μM BTZ, *NGLY1 +/-* larvae eclose at significantly higher rates when carrying the *SEL1L^(S/P, +/+)^* (83%, p<0.0001) and *SEL1L^(S/S, +/DEL)^* (96%, p<0.0001) genotypes compared to the *SEL1L^(S/S, +/+)^* genotype (59%). (B) When treated with 1μM BTZ, adult *NGLY1+/-* flies with the *SEL1L^(S/P, +/+)^* (p<0.0001) and *SEL1L^(S/S, +/DEL)^* (p<0.0001) genotypes live longer than the *SEL1L^(S/S, +/+)^*. Cox proportional hazard regression analysis. (C) There is no significant difference in survival between adult *NGLY1 +/-* flies treated with DMSO regardless of *SEL1L* genotype.

To understand if the *SEL1L* variants impact NRF1 signaling in adult flies, we treated heterozygous *NGLY1* null adult flies with bortezomib or DMSO and observed their long-term, adult survival. The adult flies were raised on DMSO during their larval stages and were either maintained on DMSO or BTZ food. *SEL1L^(S/S, +/+)^* adult flies showed significantly decreased survival time on BTZ compared to the *SEL1L^(S/P, +/+)^* (p<0.0001) and the *SEL1L^(S/S, +/DEL)^* (p<0.0001) genotypes (Fig 3B and S3 Data). The increased survival times of the bortezomib-treated *SEL1L^(S/P, +/+)^* and *SEL1L^(S/S, +/DEL)^* genotypes indicate that these variants are protective against proteasome inhibition during adulthood. When treated with DMSO, heterozygous *NGLY1* null adult flies displayed no significant differences in long-term survival, regardless of *SEL1L* variant genotypes (Fig 3C and S3 Data). These experiments demonstrate that the sensitivity to proteasome inhibition in NGLY1 deficiency is modified by these *SEL1L* variants, both during larval development and in adulthood. These results suggest that these *SEL1L* variants impact proteasome sensitivity in an NGLY1-dependent manner.

### SEL1L effects on NRF1 signaling and proteasome gene expression

In response to proteasome inhibition, NRF1 upregulates proteasome genes (21). To test for changes in NRF1 signaling in our *SEL1L* variant flies, we examined the expression of several proteasomal subunit genes: *prosalpha6*, *prosalpha3*, *prosbeta2*, *prosbeta4*, and *prosbeta5*. These five genes were among other proteasome genes that we previously identified as downregulated in our NGLY1 deficiency fly model (7). We treated flies with BTZ to induce NRF1 signaling. When we compared expression of the proteasome genes, in whole flies, between *NGLY1* wildtype and heterozygous *NGLY1* null adults, we mostly observed no differences in gene expression within each condition and *SEL1L* genotype (S2 Fig A and S4 Data). We observed increases in proteasome gene expression with the treatment of BTZ compared to DMSO; however, we found no *SEL1L* variant specific differences in gene expression in either the *NGLY1* wildtype or heterozygous *NGLY1* null flies (S2 Fig B and S4 Data). While this is unexpected in light of the previous phenotypic data we present, it is possible that there are specific effects in different tissues that are missed when we examine expression in whole flies. To date, it is unknown which tissues contribute to the lethality in NGLY1 flies and more work is needed to determine which tissues are most impacted by proteasome inhibition.

### *SEL1L^P780^* and *SEL1L^Δ806-809^* variants enhance ERAD in an ER stress model

SEL1L is an integral component of the Hrd1 ERAD complex, which retrotranslocates misfolded proteins from the ER lumen and ubiquitinates them for degradation by the proteasome (10,13,15). Although its role in ERAD remains unclear, NGLY1 has been shown to physically interact with proteins in the ERAD complex, including Derlin-1 and VCP (25,26). The deglycosylation of ERAD substrates by NGLY1 is thought to prepare misfolded glycoproteins for proteasomal degradation (5,27,28).

We crossed a *Drosophila* eye model of ER stress with the *SEL1L* strains to determine the variant-specific effects on ERAD. In this model, a transgene carries an eye-specific GAL4 (*GMR-GAL4*) that drives the overexpression of a mutant misfolded rhodopsin protein (encoded by *UAS- Rh1^G69D^*) that constitutively misfolds and induces degeneration in the developing larval eye disc (29–32). The misfolded rhodopsin protein leads to chronic ER stress in the eye disc, cell death, and a rough eye phenotype in adult flies. These eyes are also significantly smaller than wildtype eyes. This model is sensitive to changes in ERAD function and increased ERAD function is protective against eye degeneration (29). In larval eye discs, overexpressing Hrd1, the ERAD protein essential for transporting misfolded proteins out of the ER, nearly completely restored the normal appearance of the eye (29). Increasing Hrd1 levels enhances ERAD, helping to clear misfolded rhodopsin, which in turn prevents ER stress and protects against degeneration. Conversely, knockdown of ERAD components in this model leads to increased eye degeneration (29). Because SEL1L is a component of the Hrd1 ERAD complex, modifying SEL1L should similarly affect phenotypes in the ER stress eye model.

We hypothesized that losing SEL1L would decrease ERAD and enhance the eye degeneration phenotype, leading to a smaller eye size. We expressed *SEL1L RNAi* in the eye discs to knockdown *SEL1L* in the ER stress model (and NGLY1 wildtype) and observed that the eyes were significantly smaller than in the flies without knockdown of *SEL1L* (p= 0.0016) (Fig 4A and S5 Data). The reduction in eye size observed in this model following *SEL1L* knockdown is expected, given SEL1L’s established role in ERAD function. We next crossed the different *SEL1L* variants onto this model (and NGLY1 wildtype) to investigate their effects on eye size. Because the ER stress model strain carries the common *SEL1L^S780^* allele, crossing our *SEL1L* variants onto this model provided the same *SEL1L* genotypes as previously described. We observed an increase in eye size of the *SEL1L^(S/P, +/+)^* (p<0.0001) and *SEL1L^(S/S, +/DEL)^* (p<0.0001) genotypes compared to the SEL1L*^(S/S, +/+)^* flies (Fig 4B and S5 Data), opposite of what we observed with the RNAi knockdown experiment. Given that previous studies show that improvement in ERAD function increases eye size, this result suggests that the *SEL1L^P780^* and *SEL1L^Δ806-809^* alleles improve ERAD function in an NGLY1 wildtype background.

**Fig 4.**
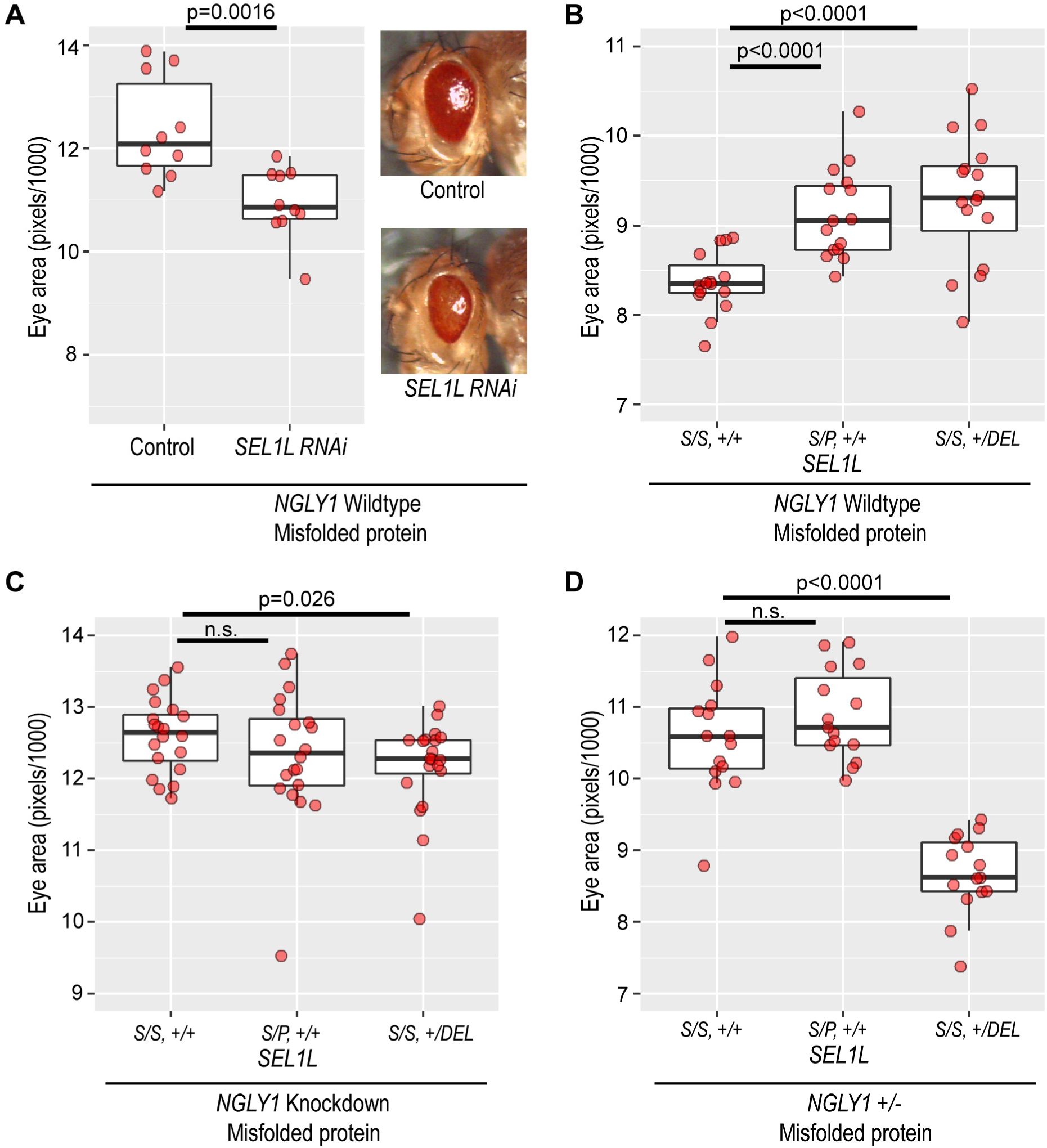
SEL1L variants enhance ERAD in an NGLY1-dependent manner. (A) Knockdown of *SEL1L* reduces eye size in an ER stress model expressing a misfolded protein in the eye (Eye size of Control 12.38 ± 0.93, *SEL1L* RNAi 10.93 ± 0.65, p=0.0016). (B) In the ER stress model, with an *NGLY1* wildtype background, *SEL1L^(S/P, +/+)^* (9.13 ± 0.49, p<0.0001) and *SEL1L^(S/S, +/DEL)^* (9.29 ± 0.69, p < 0.0001) genotypes significantly increase eye size compared to the *SEL1L^(S/S, +/+)^* genotype (8.36 ± 0.33). (C) Eye-specific knockdown of NGLY1 in the ER stress model results in no difference in eye size between the *SEL1L^(S/S, +/+)^* and *SEL1L^(S/P, +/+)^* genotypes. The *SEL1L^(S/S, +/DEL)^* genotype shows a significantly smaller eye size (p <0.05). (Eye size: *SEL1L^(S/S, +/+)^* 12.60 ± 0.50, *SEL1L^(S/P, +/+)^* 12.34 ± 0.89, and *SEL1L^(S/S, +/DEL)^* 12.16 ± 0.65). (D) Heterozygous loss of *NGLY1* in the ER stress model results in no difference in eye size between the *SEL1L^(S/S, +/+)^* and *SEL1L^(S/P, +/+)^* genotypes. The *SEL1L^(S/S, +/DEL)^* genotype shows a significantly smaller eye size (p <0.0001). (Eye size: *SEL1L^(S/S, +/+)^* 10.58 ± 0.76, *SEL1L^(S/P, +/+)^* 10.88 ± 0.61, and *SEL1L^(S/S, +/DEL)^* 8.67 ± 0.54).

We next tested whether ERAD is still improved by the *SEL1L^P780^* and *SEL1L^Δ806-809^* alleles when NGLY1 activity is reduced. When we knockdown *NGLY1* in the eye of this ER stress model, eye size is no longer increased in the *SEL1L^(S/P, +/+)^* and *SEL1L^(S/S, +/DEL)^* genotypes compared to the *SEL1L^(S/S, +/+)^* (Fig 4C and S5 Data). This result suggests that the improvement in ERAD associated with the *SEL1L^(S/P, +/+)^* and *SEL1L^(S/S, +/DEL)^* genotypes is dependent on NGLY1 activity. Heterozygous *NGLY1* null flies also showed no improvement in ERAD in the *SEL1L^(S/P, +/+)^* and *SEL1L^(S/S, +/DEL)^* genotypes compared to the *SEL1L^(S/S, +/+)^* genotype (Fig 4D and S5 Data). Interestingly, with the loss of NGLY1, the *SEL1L^(S/S, +/DEL)^* genotype had smaller eyes than the *SEL1L^(S/S, +/+)^* genotype in both the *NGLY1* knockdown (p<0.05) and heterozygous *NGLY1* null (p<0.0001). We conclude that the *SEL1L* variants improve ERAD in an NGLY1-dependent manner, further supporting the role of NGLY1 in ERAD. In the absence or reduction of NGLY1, the *SEL1L* variants do not increase ERAD in this model. We also conclude that the increased survival observed with *SEL1L^(S/P, +/+)^* and *SEL1L^(S/S, +/DEL)^* variants in the NGLY1 deficiency *Drosophila* model are likely not attributed to improvements in ERAD of misfolded proteins.

## DISCUSSION

In this study, we sought to understand how the SEL1L protein-coding variants identified in our *Drosophila* genetic screen affect NGLY1 deficiency lethality by investigating the ERAD and NRF1 signaling pathways. We conclude that the *SEL1L* variants are increasing survival in our NGLY1 deficiency models through the NRF1 signaling pathway. We observed that the *SEL1L^P780^* and *SEL1L^Δ806-809^* variants provide protection against proteasome inhibition during both larval development and in adult flies when NGLY1 was reduced. We propose that our SEL1L variants are more efficiently removing NRF1 from the ER, thereby increasing NRF1 activation by the remaining NGLY1 and resisting proteasome inhibition (Fig 5).

**Fig 5.**
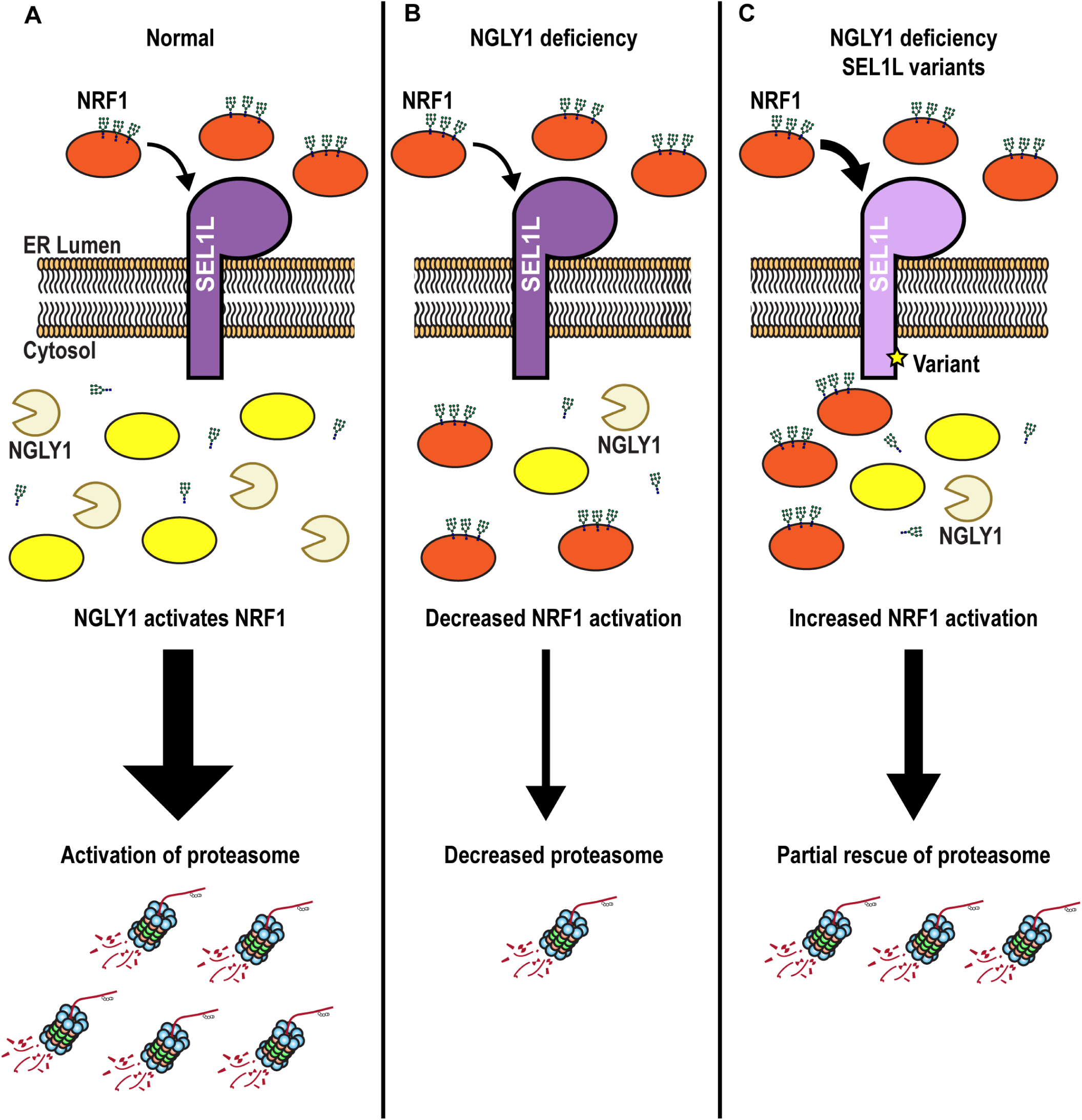
Proposed model of how SEL1L variants interact with NGLY1. (A) Under normal conditions, inactive, glycosylated NRF1 (orange oval) is removed from the ER lumen by ERAD (SEL1L) where it can be deglycosylated and activated (yellow oval) by NGLY1 and elicit a robust proteasome response. (B) In NGLY1 deficiency, less NGLY1 is available to activate NRF1, resulting in a decrease of proteasome response. (C) In NGLY1 deficiency with the *SEL1L^(S/P, +/+)^* and *SEL1L^(S/S, +/DEL)^* variants, enhanced ERAD results in an increased amount of NRF1 removed from the ER. This allows for increased NRF1 activation and increased proteasome response, despite reduced NGLY1.

The results from our ERAD experiments suggest that the SEL1L variants enhance ERAD in an NGLY1-dependent manner. While ERAD is a separate pathway from NRF1 signaling, the pathways are inextricably linked. The Lehrbach and Ruvkin NRF1 screen identified loss-of-function mutations in *SEL1L* that decreased NRF1 signaling and previous work has shown that ERAD complex components are responsible for NRF1 removal from the ER (17–19,33). If ERAD is enhanced by *SEL1L^P780^* and *SEL1L^Δ806-809^* variants on an NGLY1 deficiency background, we hypothesize that, in these genotypes, more NRF1 can be translocated from the ER. This increased NRF1 removal from the ER is likely compensating for the reduction in NGLY1 and elicits a more robust proteasome bounce-back response, despite the reduction in NGLY1 (Fig 5).

These results point to ERAD and NRF1 signaling as being potential therapeutic targets for NGLY1 deficiency patients. While there are still no therapeutic treatments for NGLY1 deficiency patients, there are two ongoing clinical trials, including an intracerebroventricular (ICV) *NGLY1* gene replacement therapy (34) and a GlcNac supplementation trial to treat alacrima, or reduced tear production (35). Further study is needed to investigate how enhancing ERAD and NRF1 signaling affect the proteasome bounce-back response in NGLY1 deficiency patients who already suffer from proteasome dysfunction (36). Most current therapeutics that target ERAD only aim to decrease its function (37–39), however, enhancing ERAD is possible in cells and animal models through increased expression of ERAD complex proteins (40,41). In addition to SEL1L, our previous modifier screen has also identified other genes involved in ERAD or ER function, including *TMEM259*, *TMTC2*, and *ERMP1* (9). Interestingly, for both *TMCT2* and *ERMP1*, our GWAS independently hit two separate fly orthologs of these genes (9). ERAD components and *NRF1* are strong candidates for modifier genes of NGLY1 deficiency patients and may be good drug targets for further development.

## METHODS

### Fly stocks and maintenance

Stocks were maintained on standard agar-dextrose-yeast medium and standard Archon Scientific glucose fly food at 25°C on a 12-h light/dark cycle. *SEL1L* CRISPR strains were created by WellGenetics, Inc (www.wellgenetics.com) and alleles were verified by Sanger sequencing (S3 Fig). The following stocks were obtained from the Bloomington *Drosophila* Stock Center (Bloomington, IN): Tubulin-GAL4 (BDSC #5138), UAS-pngl-RNAi (BDSC #54853), attp2 (BDSC #36303), and attp40 (BDSC #36304). The *Tubulin*-GAL80 strain was provided by Dr. Carl Thummel (University of Utah). The *SEL1L-RNAi* strain (v1161) was obtained from the Vienna *Drosophila* Resource Center (42). The *NGLY1* null allele carries an early stop codon in the *NGLY1* gene, and was previously characterized and generously provided by Perlara, PBC (23,24). The *NGLY1* null allele is homozygous lethal and the stock is maintained with the CyO balancer.

The “ER stress model” contains GMR-GAL4 and UAS-Rh1G69D on the second chromosome and has been previously described (30–32,43). The endogenous *SEL1L* in this strain is homozygous for the *SEL1L^P780^* allele. We backcrossed the *SEL1L^S780^* allele from our CRISPR generated *SEL1L^S780^* strain into the ER stress model for 20 generations to create the desired homozygous *SEL1L^S780^* genotype and verified genotype through sequencing.

### Proteasome sensitivity larval size assay

Proteasome sensitivity larval size assays were performed as previously described (24). Standard *Drosophila* food was melted and cooled to 60°C prior to the addition of DMSO or bortezomib. *NGLY1^+/-^* females were pre-mated with males from a CRISPR *SEL1L* variant strain for 24 hours. Mated females were placed on food containing DMSO and allowed to lay eggs for approximately 8 hours. Adult flies were removed and after four days of development, 3^rd^ instar larvae were transferred to vials containing food with DMSO or bortezomib. After two days on bortezomib containing food, larvae were genotyped using the presence of GFP. The *CyO* balancer also carries a GFP marker. Larvae were confirmed to lack the balancer chromosome by checking that they were also GFP negative. Larvae were imaged at 2.5X magnification using a Leica EC3 camera. Larval size was quantified using ImageJ as previously described (23,24).

### Survival assays

NGLY1 knockdown (KD) eclosion survival: Virgin females from the CRISPR *SEL1L* variant strains were fed yeast overnight and then crossed with males from the donor strain UAS- NGLY1RNAi/Cyo,Tubulin-GAL80; Tubulin-GAL4/TM3,Sb. Progeny were collected and scored for the four balancer classes: CyO, Sb, double balanced, or no balancers, with the no balancer flies being the NGLY1 KD. This cross should produce the expected 1:1:1:1 ratio of the four genotypes. Given that there is always a very low level of lethality associated with each balancer, the largest balancer class was considered the closest to the expected number. We scored at least 200 flies per cross. Males and females were combined for a single count. To calculate the proportion of NGLY1 KD flies by generating a ratio of NGLY1 knockdown/largest balancer class as previously described (9). For long-term survival, flies were collected, placed in a vial and flipped into fresh food every 2-3 days. Lifespan was measured as days post-eclosion. Vials were checked daily for dead flies and recorded.

### Eye imaging and quantification

Adult female flies aged 3-5 days were collected under CO2 anesthesia then frozen at −80°C for later imaging. Eyes were imaged at 3x magnification using the Leica EC3 Camera. Eye area was measured as previously described (30–32,44)

### Proteasome gene RT-qPCR

Changes in proteasome subunit gene expression were measured using RT-qPCR. Flies with appropriate *SEL1L* and *NGLY1* genotypes were crossed as previously described. 24 hours after eclosion, male flies were collected and placed on either 0.2% DMSO or 3μM BTZ for 12 hours or 24 hours. Immediately after drug treatments, RNA was extracted from 8-10 whole-body flies using a Direct-zol RNA Miniprep (Zymo Research R2061) using TRIzol Reagent (ThermoFisher Cat # 15596026) and including the DNAse step. RNA was converted to cDNA using a ProtoScript® II First Strand cDNA Synthesis Kit (NEB Cat # E6560L). RT-qPCR was performed using a QuantStudio 3 96-well 0.2 ml block instrument and PowerUp SYBR Green Master Mix (ThermoFisher Cat #A25741). If available, we used primers from the FlyPrimerBank (45) located at http://www.flyrnai.org/flyprimerbank. Other primers were designed using Primer3Plus (46) located at www.primer3plus.com. All primer sequences listed in S4 Data.

## Supporting information

S1 Data. NGLY1 ubiquitous knockdown fly counts and long-term survival.

S2 Data. Larval size assay measurements.

S3 Data. Fly counts for eclosion on BTZ and survival data.

S1 Fig. Wildtype NGLY1 larvae are unaffected by proteasome inhibition.

S2 Fig. Proteasome gene expression changes in response to BTZ.

S4 Data. qPCR results and primers used.

S5 Data. Fly eye measurements with ER stress model.

S3 Fig. Sequencing of CRISPR fly lines confirm appropriate SEL1L variants.

## Supporting Information

**S1 Data.** *NGLY1* ubiquitous knockdown fly counts and long-term survival.

**S2 Data.** Larval size assay measurements.

**S3 Data.** Fly counts for eclosion on BTZ and survival data.

**S1 Fig.** Wildtype *NGLY1* larvae are unaffected by proteasome inhibition.

**S2 Fig.** Proteasome gene expression changes in response to BTZ.

**S4 Data.** qPCR results and primers used.

**S5 Data.** Fly eye measurements with ER stress model.

**S3 Fig.** Sequencing of CRISPR fly lines confirm appropriate *SEL1L* variants.

## REFERENCES

1. Freeze HH. Understanding Human Glycosylation Disorders: Biochemistry Leads the Charge. J Biol Chem. 2013 Mar 8;288(10):6936–45.

2. Enns GM, Shashi V, Bainbridge M, Gambello MJ, Zahir FR, Bast T, et al. Mutations in NGLY1 cause an inherited disorder of the endoplasmic reticulum–associated degradation pathway. Genetics in Medicine. 2014 Oct;16(10):751–8.

3. Suzuki T, Huang C, Fujihira H. The cytoplasmic peptide:N-glycanase (NGLY1); structure, expression and cellular functions. Gene. 2016 Feb 10;577(1):1–7.

4. Hosomi A, Fujita M, Tomioka A, Kaji H, Suzuki T. Identification of PNGase-dependent ERAD substrates in Saccharomyces cerevisiae. Biochem J. 2016 Oct 1;473(19):3001–12.

5. Hirsch C, Blom D, Ploegh HL. A role for N-glycanase in the cytosolic turnover of glycoproteins. EMBO J. 2003 Mar 3;22(5):1036–46.

6. Kario E, Tirosh B, Ploegh HL, Navon A. N-Linked Glycosylation Does Not Impair Proteasomal Degradation but Affects Class I Major Histocompatibility Complex Presentation. J Biol Chem. 2008 Jan 4;283(1):244–54.

7. Owings KG, Lowry JB, Bi Y, Might M, Chow CY. Transcriptome and functional analysis in a Drosophila model of NGLY1 deficiency provides insight into therapeutic approaches. Hum Mol Genet. 2018 Mar 15;27(6):1055–66.

8. Caglayan AO, Comu S, Baranoski JF, Parman Y, Kaymakçalan H, Akgumus GT, et al. NGLY1 Mutation Causes Neuromotor Impairment, Intellectual Disability, and Neuropathy. Eur J Med Genet. 2015 Jan;58(1):39–43.

9. Talsness DM, Owings KG, Coelho E, Mercenne G, Pleinis JM, Partha R, et al. A Drosophila screen identifies NKCC1 as a modifier of NGLY1 deficiency. Bellen HJ, Wittkopp PJ, Tiemeyer M, editors. eLife. 2020 Dec 14;9:e57831.

10. Sun S, Shi G, Han X, Francisco AB, Ji Y, Mendonça N, et al. Sel1L is indispensable for mammalian endoplasmic reticulum-associated degradation, endoplasmic reticulum homeostasis, and survival. PNAS. 2014 Feb 4;111(5):E582–91.

11. Mehnert M, Sommermeyer F, Berger M, Kumar Lakshmipathy S, Gauss R, Aebi M, et al. The interplay of Hrd3 and the molecular chaperone system ensures efficient degradation of malfolded secretory proteins. MBoC. 2015 Jan 15;26(2):185–94.

12. Jeong H, Sim HJ, Song EK, Lee H, Ha SC, Jun Y, et al. Crystal structure of SEL1L: Insight into the roles of SLR motifs in ERAD pathway. Sci Rep. 2016 Feb 9;6(1):20261.

13. Mueller B, Lilley BN, Ploegh HL. SEL1L, the homologue of yeast Hrd3p, is involved in protein dislocation from the mammalian ER. J Cell Biol. 2006 Oct 23;175(2):261–70.

14. Mueller B, Klemm EJ, Spooner E, Claessen JH, Ploegh HL. SEL1L nucleates a protein complex required for dislocation of misfolded glycoproteins. PNAS. 2008 Aug 26;105(34):12325–30.

15. Iida Y, Fujimori T, Okawa K, Nagata K, Wada I, Hosokawa N. SEL1L protein critically determines the stability of the HRD1-SEL1L endoplasmic reticulum-associated degradation (ERAD) complex to optimize the degradation kinetics of ERAD substrates. J Biol Chem. 2011 May 13;286(19):16929–39.

16. Hosokawa N, Wada I. Association of the SEL1L protein transmembrane domain with HRD1 ubiquitin ligase regulates ERAD-L. The FEBS Journal. 2016;283(1):157–72.

17. Lehrbach NJ, Ruvkun G. Proteasome dysfunction triggers activation of SKN-1A/Nrf1 by the aspartic protease DDI-1. Dillin A, editor. eLife. 2016 Aug 16;5:e17721.

18. Tomlin FM, Gerling-Driessen UIM, Liu YC, Flynn RA, Vangala JR, Lentz CS, et al. Inhibition of NGLY1 Inactivates the Transcription Factor Nrf1 and Potentiates Proteasome Inhibitor Cytotoxicity. ACS Cent Sci. 2017 Nov 22;3(11):1143–55.

19. Radhakrishnan SK, den Besten W, Deshaies RJ. p97-dependent retrotranslocation and proteolytic processing govern formation of active Nrf1 upon proteasome inhibition. Brown MS, editor. eLife. 2014 Jan 21;3:e01856.

20. Koizumi S, Irie T, Hirayama S, Sakurai Y, Yashiroda H, Naguro I, et al. The aspartyl protease DDI2 activates Nrf1 to compensate for proteasome dysfunction. Dikic I, editor. eLife. 2016 Aug 16;5:e18357.

21. Radhakrishnan SK, Lee CS, Young P, Beskow A, Chan JY, Deshaies RJ. Transcription Factor Nrf1 Mediates the Proteasome Recovery Pathway after Proteasome Inhibition in Mammalian Cells. Molecular Cell. 2010 Apr 9;38(1):17–28.

22. Mackay TFC, Richards S, Stone EA, Barbadilla A, Ayroles JF, Zhu D, et al. The Drosophila melanogaster Genetic Reference Panel. Nature. 2012 Feb;482(7384):173–8.

23. Rodriguez TP, Mast JD, Hartl T, Lee T, Sand P, Perlstein EO. Defects in the Neuroendocrine Axis Contribute to Global Development Delay in a Drosophila Model of NGLY1 Deficiency. G3: Genes, Genomes, Genetics. 2018 Jul 1;8(7):2193–204.

24. Hope KA, Berman AR, Peterson RT, Chow CY. An in vivo drug repurposing screen and transcriptional analyses reveals the serotonin pathway and GSK3 as major therapeutic targets for NGLY1 deficiency. PLOS Genetics. 2022 Feb 6;18(6):e1010228.

25. Katiyar S, Joshi S, Lennarz WJ. The retrotranslocation protein Derlin-1 binds peptide:N- glycanase to the endoplasmic reticulum. Molecular Biology of the Cell. 2005;16(10):4584–94.

26. McNEILL H, Knebel A, Arthur JSC, Cuenda A, Cohen P. A novel UBA and UBX domain protein that binds polyubiquitin and VCP and is a substrate for SAPKs. Biochem J. 2004 Dec 1;384(2):391–400.

27. Bebök Z, Mazzochi C, King SA, Hong JS, Sorscher EJ. The mechanism underlying cystic fibrosis transmembrane conductance regulator transport from the endoplasmic reticulum to the proteasome includes Sec61beta and a cytosolic, deglycosylated intermediary. J Biol Chem. 1998 Nov 6;273(45):29873–8.

28. Misaghi S, Pacold ME, Blom D, Ploegh HL, Korbel GA. Using a Small Molecule Inhibitor of Peptide: N-Glycanase to Probe Its Role in Glycoprotein Turnover. Chemistry & Biology. 2004 Dec 1;11(12):1677–87.

29. Kang MJ, Ryoo HD. Suppression of retinal degeneration in Drosophila by stimulation of ER- associated degradation. Proc Natl Acad Sci U S A. 2009 Oct 6;106(40):17043–8.

30. Chow CY, Kelsey KJP, Wolfner MF, Clark AG. Candidate genetic modifiers of retinitis pigmentosa identified by exploiting natural variation in Drosophila. Hum Mol Genet. 2016 Feb 15;25(4):651–9.

31. Palu RAS, Chow CY. Baldspot/ELOVL6 is a conserved modifier of disease and the ER stress response. Lin JH, editor. PLoS Genet. 2018 Aug 6;14(8):e1007557.

32. Palu RAS, Dalton HM, Chow CY. Decoupling of Apoptosis from Activation of the ER Stress Response by the Drosophila Metallopeptidase superdeath. Genetics. 2020 Apr 1;214(4):913– 25.

33. Steffen J, Seeger M, Koch A, Krüger E. Proteasomal Degradation Is Transcriptionally Controlled by TCF11 via an ERAD-Dependent Feedback Loop. Molecular Cell. 2010 Oct 8;40(1):147–58.

34. Grace Science, LLC. A Phase 1/2/3 Open-label, Single Arm, Dose-finding Study to Investigate Long-term Safety, Tolerability and Efficacy of GS-100, an Adeno-associated Virus Serotype 9 (AAV9) Vector-mediated Gene Transfer of Human NGLY1, in Patients With NGLY1 Deficiency [Internet]. clinicaltrials.gov; 2024 Jun [cited 2025 Jan 31]. Report No.: NCT06199531. Available from: https://clinicaltrials.gov/study/NCT06199531

35. Morava-Kozicz E. A Phase II Randomized, Multicenter, Double-Blind, Placebo-Controlled Study Evaluating Effect Of GlcNAc On Tear Production In Individuals With NGLY1-CDDG [Internet]. clinicaltrials.gov; 2025 Jan [cited 2025 Jan 31]. Report No.: NCT05402345. Available from: https://clinicaltrials.gov/study/NCT05402345

36. Yoshida Y, Asahina M, Murakami A, Kawawaki J, Yoshida M, Fujinawa R, et al. Loss of peptide:N-glycanase causes proteasome dysfunction mediated by a sugar-recognizing ubiquitin ligase. PNAS [Internet]. 2021 Jul 6 [cited 2021 Jul 6];118(27). Available from: https://www.pnas.org/content/118/27/e2102902118

37. Li X, Zhang K, Li Z. Unfolded protein response in cancer: the Physician’s perspective. Journal of Hematology & Oncology. 2011 Feb 23;4(1):8.

38. Harbut MB, Patel BA, Yeung BKS, McNamara CW, Bright AT, Ballard J, et al. Targeting the ERAD pathway via inhibition of signal peptide peptidase for antiparasitic therapeutic design. Proc Natl Acad Sci U S A. 2012 Dec 26;109(52):21486–91.

39. Kim H, Bhattacharya A, Qi L. Endoplasmic reticulum quality control in cancer: Friend or foe. Seminars in Cancer Biology. 2015 Aug 1;33:25–33.

40. Belmont PJ, Chen WJ, San Pedro MN, Thuerauf DJ, Lowe NG, Gude N, et al. Roles for ER- associated Degradation (ERAD) and the Novel ER Stress Response Gene, Derlin-3, in the Ischemic Heart. Circ Res. 2010 Feb 5;106(2):307–16.

41. Doroudgar S, Völkers M, Thuerauf DJ, Khan M, Mohsin S, Respress JL, et al. Hrd1 and ER-Associated Protein Degradation, ERAD, Are Critical Elements of the Adaptive ER Stress Response in Cardiac Myocytes. Circ Res. 2015 Aug 28;117(6):536–46.

42. Dietzl G, Chen D, Schnorrer F, Su KC, Barinova Y, Fellner M, et al. A genome-wide transgenic RNAi library for conditional gene inactivation in Drosophila. Nature. 2007 Jul;448(7150):151– 6.

43. Ryoo HD, Domingos PM, Kang MJ, Steller H. Unfolded protein response in a Drosophila model for retinal degeneration. The EMBO Journal [Internet]. 2006 Dec 14 [cited 2024 Dec 6]; Available from: https://www.embopress.org/doi/10.1038/sj.emboj.7601477

44. Dalton HM, Viswanatha R, Jr RB, Zuno JS, Berman AR, Rushforth R, et al. A genome-wide CRISPR screen identifies DPM1 as a modifier of DPAGT1 deficiency and ER stress. PLOS Genetics. 2022 Sep 27;18(9):e1010430.

45. Hu Y, Sopko R, Foos M, Kelley C, Flockhart I, Ammeux N, et al. FlyPrimerBank: An Online Database for Drosophila melanogaster Gene Expression Analysis and Knockdown Evaluation of RNAi Reagents. G3 Genes|Genomes|Genetics. 2013 Sep 1;3(9):1607–16.

46. Untergasser A, Cutcutache I, Koressaar T, Ye J, Faircloth BC, Remm M, et al. Primer3--new capabilities and interfaces. Nucleic Acids Res. 2012 Aug;40(15):e115.

