## Supplementary figures and images for "Natural *SEL1L* variants modify ERAD, proteasome function, and survival in a *Drosophila* model of NGLY1 deficiency"

### S1 Fig. Wildtype NGLY1 larvae are unaffected by proteasome inhibition.

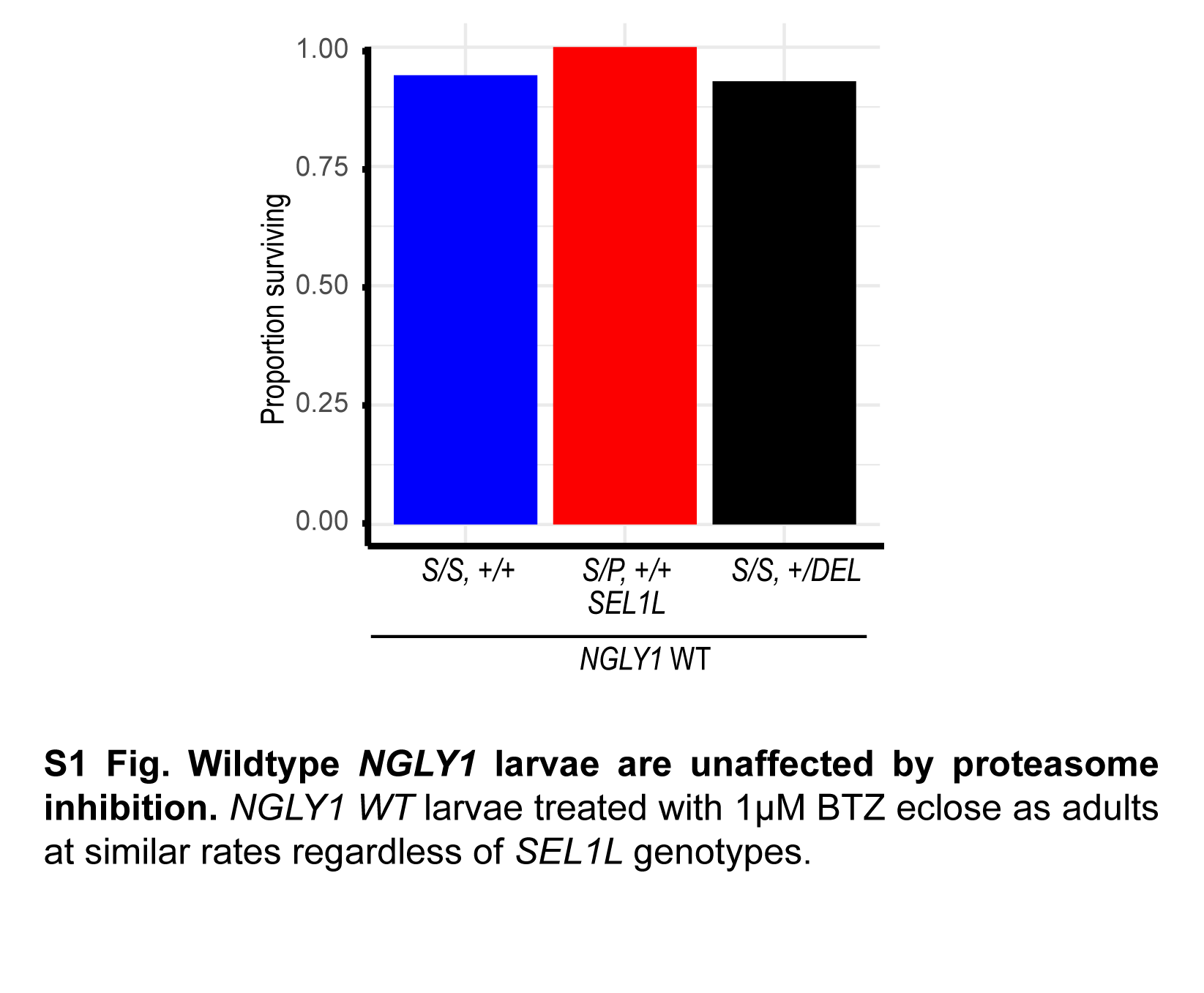

### S2 Fig. Proteasome gene expression changes in response to BTZ.

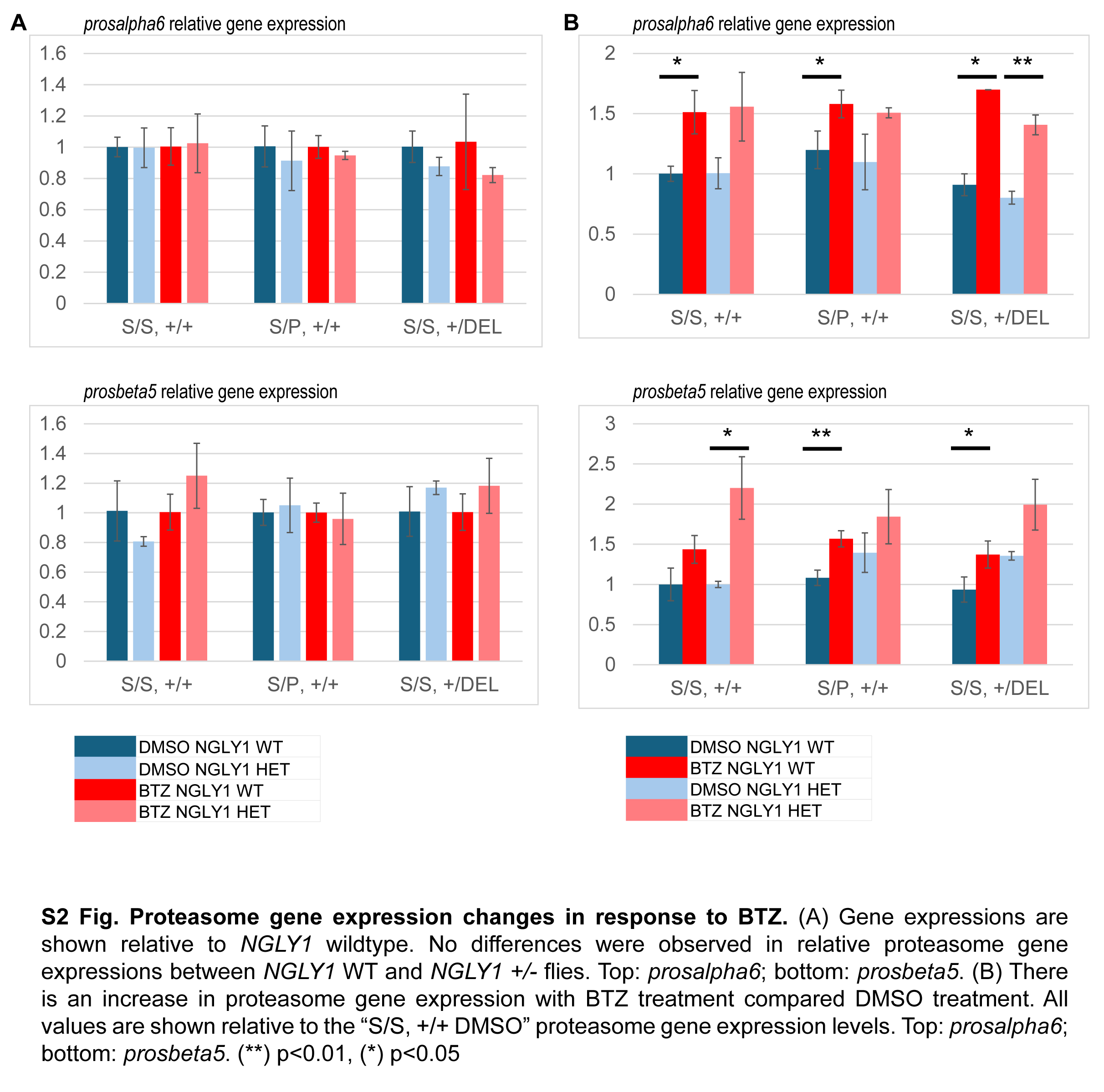

### S3 Fig. Sequencing of CRISPR fly lines confirm appropriate SEL1L variants.

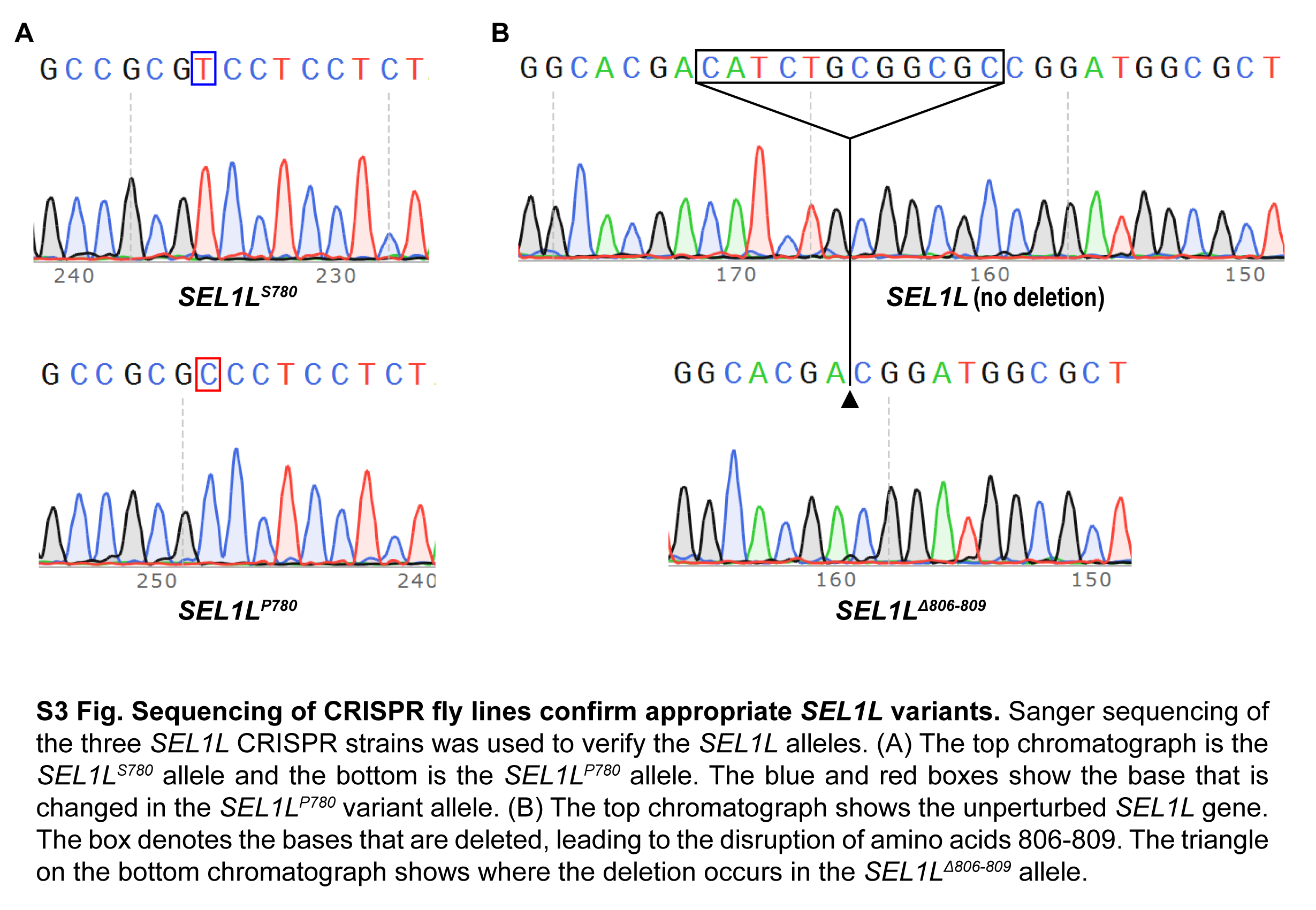
